# Astatine-211 injury induces an immunostimulatory melanoma-cell state and promotes systemic antitumor immunity

**DOI:** 10.64898/2026.10.22.760895

**Authors:** Koki Mayusumi, Soichiro Magara, Xuhao Huang, Kazuko Kaneda-Nakashima, Kazuhiro Ooe, Masashi Murakami, Atsushi Toyoshima, Hiromitsu Haba, Xiaojie Yin, Koichi Fukase, Atsushi Shimoyama, Kazuya Kabayama

## Abstract

Targeted alpha therapy is generally evaluated by absorbed dose and direct tumor-cell killing, but whether alpha-particle-injured tumor cells can induce systemic antitumor immunity remains poorly defined. We investigated the immunogenic consequences of astatine-211 (²¹¹At) injury in a prophylactic melanoma vaccination–challenge model. B16F10 cells were exposed ex vivo to ²¹¹At, etoposide, or ultraviolet irradiation under conditions that reduced viable-cell counts at 24 h to approximately 40% of untreated controls. Injured cells were administered subcutaneously to syngeneic C57BL/6J mice before intravenous challenge with viable B16F10 cells. Vaccination with cells exposed to 1 MBq/mL ²¹¹At suppressed pulmonary metastasis, reduced regulatory T-cell proportions, and elicited tumor-reactive interferon-γ responses and antibodies. Transcriptomic profiling revealed enrichment of type I interferon, inflammatory, and chemokine-related signatures, together with suppression of cell-cycle, translation, and metabolic programs. At the cellular level, ²¹¹At injury was accompanied by γH2AX accumulation, G2/M arrest, increased reactive oxygen species and cytosolic double-stranded DNA, and senescence-like changes. These features arose without consistent increases in classical damage-associated molecular patterns or prominent induction of senescence-associated secretory phenotype factors at 1 MBq/mL. In vivo pharmacologic inhibition of stimulator of interferon genes (STING) signaling attenuated the antimetastatic effect of vaccination. Together, these findings demonstrate that ²¹¹At injury can generate an immunostimulatory tumor-cell state characterized by DNA damage and altered inflammatory signaling. STING contributes to the resulting immune protection, highlighting immunogenic tumor-cell conversion as a consideration in the design of targeted alpha therapies.

## Introduction

Targeted alpha therapy has attracted increasing attention as a cancer treatment, and radionuclides such as ²²³Ra (half-life, 11 d) and ²²□Ac (half-life, 9.9 d) have been extensively investigated (1–5). Among alpha-particle-emitting radionuclides, astatine-211 (²¹¹At) is of particular interest. Its 7.2-h physical half-life permits radiochemical production, purification and short biological procedures while delivering radiation over a relatively brief period. ²¹¹At decays either by alpha emission or by electron capture; the electron-capture branch produces short-lived ²¹¹Po, which rapidly undergoes alpha decay. The high linear energy transfer (LET) of the resulting alpha particles produces clustered DNA damage over a short tissue range. ²¹¹At can also be produced by cyclotron irradiation of bismuth and incorporated into small molecules, peptides and antibodies (6–8).

Clinical development of ²¹¹At radiopharmaceuticals remains less mature than that of beta-particle emitters or ²²³Ra, but human studies have established the feasibility of ²¹¹At radioimmunotherapy and more recent ²¹¹At-based targeted alpha therapy (9–12). These developments make it important to understand not only direct cytotoxicity but also the biological state adopted by tumor cells after ²¹¹At exposure.

We selected ²¹¹At for two principal reasons. First, high-LET alpha radiation is expected to generate densely clustered, difficult-to-repair DNA lesions that can lead to persistent proliferative arrest, micronuclei or cytosolic DNA (1, 13, 14). Second, because ²¹¹At radiopharmaceuticals are entering clinical development, the possibility that ²¹¹At-damaged tumor cells become immunostimulatory is directly relevant to treatment design and combination strategies with immunotherapy. Alpha irradiation can induce antitumor immunity, and targeted alpha therapy may cooperate with immune-checkpoint inhibitors (ICIs) (15, 16), but the immunological state induced specifically by ²¹¹At remains incompletely defined.

Classical immunogenic cell death is associated with coordinated calreticulin exposure, adenosine triphosphate (ATP) release, high-mobility group box 1 (HMGB1) release and heat-shock-protein-associated signals (17–20). However, viable tumor cells bearing persistent DNA damage can also promote immunity (21). Cytosolic dsDNA is sensed through the cGAS–STING pathway, leading to type I interferon (IFN) and chemokine responses; in irradiated tumors, STING-dependent DNA sensing supports cross-priming and CD8□ T-cell-dependent antitumor immunity (13, 14). Persistent DNA damage is also linked to therapy-induced senescence, cytosolic chromatin fragments and inflammatory secretory programs (22, 23).

Using an ex vivo tumor-cell injury model and a syngeneic pulmonary-metastasis model, we investigated whether ²¹¹At injury enables poorly immunogenic B16F10 melanoma cells to induce systemic antitumor immunity. We integrated in vivo and in vitro immune assays with characterization of proliferative arrest, DNA damage, senescence-like features, DAMPs, cytosolic dsDNA and transcriptional programs, and tested the functional contribution of STING signaling pharmacologically.

## Materials and Methods

### Cell culture

Mouse melanoma B16F10 cells (RIKEN BRC, RCB2630, RRID: CVCL_0159), mouse Lewis lung carcinoma (LLC) cells (Procell, CL-0140, RRID: CVCL_4358) and mouse macrophage-like RAW264.7 cells (ATCC, TIB-71, RRID: CVCL_0493) were used. LLC cells were kindly provided by Professor Fei Yu (Tongji University), and RAW264.7 cells were kindly provided by Dr. Hirotaka Kanoh (Keio University). All cells tested negative for *Mycoplasma*. LLC cells were cultured in RPMI-1640 medium (FUJIFILM Wako Pure Chemical Corporation, 189-02025), whereas B16F10 and RAW264.7 cells were cultured in high-glucose DMEM (FUJIFILM Wako Pure Chemical Corporation, 043-30085). Media were supplemented with 10% fetal bovine serum (FBS; Corning, 35-076-CV) and 1% penicillin-streptomycin (P/S; FUJIFILM Wako Pure Chemical Corporation, 168-23191), and cells were incubated at 37 °C in 5% CO□. For ²¹¹At treatment, preparation of damaged cells and collection of conditioned medium, cells were maintained in medium containing 1% FBS and 1% P/S unless otherwise stated.

### Mice and ethics

Female C57BL/6J mice aged 5–6 weeks were purchased from Japan SLC Inc. (RRID: MGI:5488963). Animals were housed at 25 ± 1 °C under a 12-h light/dark cycle (lights on 08:00–20:00) with free access to food and water and were acclimatized for 1 week before experiments. Animal procedures were approved by the Animal Care and Use Committee of the Graduate School of Science, The University of Osaka (approval number: IRS25-009).

### Production and isolation of ²¹¹At

²¹¹At was produced using the K70 AVF cyclotron at the RIKEN RI Beam Factory (Saitama, Japan) (24) or the K140 AVF cyclotron at the Research Center for Nuclear Physics (RCNP), The University of Osaka (Osaka, Japan). Both facilities used the ²□□Bi(α,2n)²¹¹At reaction. The produced ²¹¹At was purified by dry distillation to obtain an aqueous ²¹¹At solution (25).

### Preparation of damaged B16F10 cells for vaccination

B16F10 cells were exposed ex vivo to 0.1 or 1 MBq/mL ²¹¹At, 50 μM etoposide (FUJIFILM Wako Pure Chemical Corporation, 055-08431), or 25 J/m² ultraviolet irradiation and were harvested 24 h later for vaccination. Both adherent and nonadherent cells were collected. Adherent cells were detached using PBS containing 1 mM EDTA and 1% FBS, combined with the nonadherent cell fraction, and washed two times with PBS. The cells were subsequently resuspended in PBS for administration. The interval between cell harvest and injection was kept to a minimum, and the cells were administered within 30 min of harvest.

Residual ²¹¹At activity associated with the harvested cells was measured using a 2480 WIZARD² Automatic Gamma Counter (PerkinElmer). For activity calibration, a standard aqueous ²¹¹At solution was independently measured using both the 2480 WIZARD² and a Broad Energy Germanium Detector (CANBERRA), and the resulting calibration relationship was used to convert WIZARD² measurements to activity in Bq. Measured activities were decay-corrected to the time of cell administration based on the physical decay of ²¹¹At. In the 1 MBq/mL ²¹¹At group, the residual activity at the time of administration was approximately 0.92 kBq per mouse. Trypan Blue staining results obtained 24 h after each treatment are shown in Supplementary Table S1. Absorbed dose at the cellular or host level was not determined.

### Bone-marrow isolation and differentiation into BMDCs and BMDMs

Bone-marrow cells were collected from the femurs and tibias of 7- to 9-week-old C57BL/6J mice in RPMI-1640 containing 1% FBS, and erythrocytes were lysed using RBC Lysis Buffer (BioLegend, 420301). Cells were washed and cultured for 7 days in RPMI-1640 containing 10% FBS, 1% P/S, recombinant mouse granulocyte-macrophage colony-stimulating factor (GM-CSF, 20 ng/mL; BioLegend, 576304) and IL-4 (20 ng/mL; BioLegend, 574304) to differentiate BMDCs, or in RPMI-1640 containing 10% FBS, 1% P/S and macrophage colony-stimulating factor (M-CSF, 20 ng/mL; BioLegend, 576404) for 7 days to differentiate BMDMs.

### Lung-metastasis suppression model using ²¹¹At-damaged cancer cells

Female C57BL/6J mice were injected subcutaneously with B16F10 cells damaged for 24 h (2.5 × 10□ cells in 150 μL PBS) on day 0. On day 7, viable B16F10 cells (2 × 10□ cells in 100 μL PBS) were injected via the tail vein. Mice were euthanized on day 21 and lungs were collected to evaluate pulmonary nodules. Anti-PD-1 (Bio X Cell, BE0146, RRID: AB_10949053, clone: RMP1-14) and anti-CTLA-4 (Bio X Cell, BE0164, RRID: AB_10949609, clone: 9D9) antibodies were administered intraperitoneally at 200 μg/mouse twice weekly for three doses.

### Rechallenge, immune-cell depletion and STING inhibition

For rechallenge, immune-cell-depletion and STING-inhibition experiments, B16F10 cells damaged for 24 h (1 × 10□ cells in 100 μL PBS) were injected subcutaneously on day 0, and viable B16F10 cells (2 × 10□ cells in 100 μL PBS) were injected via the tail vein on day 7. Anti-PD-1 and anti-CTLA-4 antibodies were administered intraperitoneally at 150 μg/mouse twice weekly for three doses. In the rechallenge experiment, damaged-cell-treated mice and naïve control mice were challenged intravenously with B16F10 cells (2 × 10□ cells in 100 μL PBS) on day 21; pulmonary nodules, plasma and splenocytes were analyzed on day 27. For immune-cell depletion, anti-CD4 (Bio X Cell, BP0003-1, RRID: AB_2687763, clone: GK1.5), anti-CD8 (Bio X Cell, BP0061, RRID: AB_2891359, clone: 2.43), anti-CD122 (Bio X Cell, BE0298, RRID: AB_2687820, clone: TM-Beta-1) or isotype-control antibody (Bio X Cell, BP0090, RRID: AB_2891360, clone: LTF-2) was administered intraperitoneally at 150 μg/mouse on days 5, 7, 10 and 13. For STING inhibition, C-176 (Selleck Chemicals, S6575) was dissolved in corn oil (Selleck Chemicals, S6701) and administered intraperitoneally at 750 nmol in 200 μL/mouse daily from day 0 to day 7.

### Plasma cytokine and chemokine analysis

Cytokine and chemokine profiles in mouse plasma were measured using the LEGENDplex Mouse Anti-Virus Response Panel (13-plex) (BioLegend, BL740622) and LEGENDplex Mouse Macrophage/Microglia Panel (13-plex) (BioLegend, BL740846) according to the manufacturer’s protocols. Data were analyzed with LEGENDplex Cloud-based Data Analysis Software.

### Flow-cytometric assessment of APC activation

For APC-activation experiments, B16F10 cells were treated under each condition, the medium was replaced after 2 h and cells were cultured for a further 46 h. Damaged cancer cells comprised adherent cells together with the cell pellet obtained after centrifugation of recovered culture supernatant at 1000 rpm for 3 min. The remaining supernatant was centrifuged at 2000 × g for 10 min and used as damaged-cell-conditioned medium. BMDCs, BMDMs or RAW264.7 cells (3.0 × 10□ cells) were seeded in 24-well plates 1 day before collection of damaged cancer cells. Damaged cancer cells (1.2 × 10□ cells) or conditioned medium were added for 24 h. Cells were stained with Live/Dead Fixable Lime (Invitrogen, L34990; 1:1000), blocked with anti-CD16/32 (BioLegend, 101320; 1:50) and stained with PE-Cy7 anti-mouse CD86 (BD Biosciences, 560582; 1:200) and FITC anti-mouse CD11b (BD Biosciences, 561688; 1:200). CD86 expression was assessed by flow cytometry.

### Splenic T-cell immunophenotyping

Splenocytes (1 × 10□ cells/well) were stained with Live/Dead Fixable Lime and blocked with anti-CD16/32. Surface staining used antibodies against CD44 (Thermo Fisher Scientific, 404-0441-80, RRID: AB_2925504, clone: IM7; 1:80), CD4 (Thermo Fisher Scientific, 63-0042-82, RRID: AB_2637461, clone: RM4-5; 1:160), CD62L (Thermo Fisher Scientific, 67-0621-82, RRID: AB_2717151, clone: MEL-14; 1:667), CD3 (Thermo Fisher Scientific, 53-0031-82, RRID: AB_469889, clone: 145-2C11; 1:50), CD25 (Thermo Fisher Scientific, 25-0251-82, RRID: AB_469608, clone: PC61.5; 1:160), CD8 (Thermo Fisher Scientific, 56-0081-82, RRID: AB_494005, clone: 53-6.7; 1:160) and CD45 (Thermo Fisher Scientific, 47-0451-80, RRID: AB_1548790, clone: 30-F11; 1:160), followed by intracellular FOXP3 (Thermo Fisher Scientific, 17-5773-82, RRID: AB_469457, clone: FJK-16s; 1:20) staining. Fluorescence was measured using an Attune NxT Acoustic Focusing Cytometer (Invitrogen), and data were analyzed using FlowJo version 10.10.01 (BD Biosciences). The gating strategy is shown in Supplementary Fig. S1.

### Measurement of B16F10-binding antibodies in mouse plasma

B16F10-binding plasma antibodies were measured by flow cytometry as previously described (26). B16F10 cells (3.0 × 10□) were washed with 1% BSA/PBS and incubated with mouse plasma diluted 1:50 for 1 h at 4 °C. After washing, Alexa Fluor 488 goat anti-mouse IgG (H+L) (Thermo Fisher Scientific, A11001, RRID: AB_2534069; 1:2000) was added for 1 h at 4 °C. Fluorescence intensity was measured using an Attune NxT cytometer.

### Relative WST-8 signal and LDH-release assays

B16F10 cells were seeded at 2.5 × 10³ cells/well in 96-well plates 1 day before treatment. B16F10 cells were exposed to 0.01, 0.1 or 1 MBq/mL ²¹¹At, 50 μM etoposide or 0.5 μM doxorubicin (FUJIFILM Wako Pure Chemical Corporation, 040-21521). Cell Counting Kit-8 (Dojindo, 343-07623) was used to measure WST-8 signal. Of the 200 μL culture supernatant, 100 μL was removed for the LDH assay; 10 μL CCK-8 solution was added to each well and cells were incubated for 1 h at 37 °C before absorbance at 450 nm was measured by Infinite F50 (Tecan). Membrane damage was assessed using the Cytotoxicity LDH Assay Kit-WST (Dojindo, 343-91753) according to the manufacturer’s protocol. A lysis-buffer-treated condition served as the positive control for LDH release.

### Cell-cycle analysis

B16F10 cells were seeded at 1.0 × 10□ cells/well in 6-well plates 1 day before treatment. At 48 h, cells were collected, washed and fixed overnight in 70% ethanol at 4 °C. Cells were stained with propidium iodide (50 μg/mL; Dojindo, 341-07881), RNase A (0.1 mg/mL; Thermo Fisher Scientific, EN0531) and Triton X-100 (0.05%; Nacalai Tesque, 35501-15) for 40 min at 4 °C in the dark and analyzed using an Attune NxT cytometer.

### Nucleolar morphology and SA-**β**-gal

For nucleolar imaging, B16F10 cells were seeded at 2.0 × 10□ cells/well in 24-well glass-bottom plates, fixed with 4% paraformaldehyde, permeabilized with 1% Triton X-100/PBS and stained with Nucleolus Bright Green (Dojindo, 341-09341) and Hoechst 33342 (Invitrogen, H3570). Images were acquired with a KEYENCE all-in-one fluorescence microscope (KEYENCE, BZ-X810) using a CFI Plan Apo Lambda 40× objective (NA 0.95; Nikon). Senescence-associated β-galactosidase activity was assessed with the Cellular Senescence Detection Kit - SPiDER-βGal (Dojindo, 347-09181) according to the manufacturer’s protocol.

### Microarray and GSEA

B16F10 cells were seeded at 1.0 × 10□ cells/well in 6-well plates. Total RNA was extracted from untreated cells or cells exposed to 1 MBq/mL ²¹¹At for 24 h. Samples were analyzed using the Clariom S Assay, mouse cartridge (Thermo Fisher Scientific, 902930), and arrays were scanned using a GeneChip Scanner 3000 7G (Affymetrix, 00-0210). GSEA was performed using GSEA v4.4.0 in GSEAPreranked mode with Mouse MSigDB v2026.1.Mm gene sets, including Hallmark, Reactome and Gene Ontology Biological Process (27, 28). Genes were ranked according to a metric calculated as sign(log□ fold change) × −log□□(P value), where the fold change was calculated for ²¹¹At-treated cells relative to untreated cells. Positive ranking scores therefore represented genes upregulated in ²¹¹At-treated cells, whereas negative scores represented genes downregulated in ²¹¹At-treated cells. Accordingly, positive normalized enrichment score (NES) values indicate enrichment in ²¹¹At-treated cells, whereas negative NES values indicate enrichment in untreated cells. Enrichment scores were calculated using the weighted enrichment statistic, and statistical significance was assessed using 1,000 gene-set permutations. Gene sets containing 15–500 genes were included in the analysis.

### ROS and cytosolic dsDNA

Intracellular ROS were detected using the Intracellular Reactive Oxygen Species Detection Assay Kit (Tokyo Chemical Industry, I1265). B16F10 cells were seeded in 24-well glass-bottom plates at 1.0 × 10□ cells/well; at 30 min after treatment, cells were stained according to the kit protocol and imaged using a KEYENCE microscope with a CFI Plan Fluor 20× objective (NA 0.45; Nikon). Cytosolic dsDNA was evaluated by immunofluorescence based on a reported method (29). Cells were fixed 24 or 48 h after treatment, permeabilized with 0.1% Tween-20 and 0.01% Triton X-100, incubated with anti-dsDNA antibody (Abcam, ab27156, RRID: AB_470907, clone: 35I9 DNA; 1:1000) overnight at 4 °C and then with Alexa Fluor 488 goat anti-rabbit IgG (H+L) (Abcam, ab150077, RRID: AB_2630356; 1:200). Nuclei were stained with Hoechst 33342. Cytosolic dsDNA foci were quantified using Fiji (30).

### Splenocyte preparation and ELISPOT

At the endpoint, spleens were passed through a 100-μm cell strainer (Corning, 431752), erythrocytes were lysed with RBC Lysis Buffer and splenocytes were resuspended in T-cell culture medium consisting of RPMI-1640, 10% FBS, 1% P/S, 1 mM sodium pyruvate (Gibco, 11360070), 50 μM 2-mercaptoethanol (FUJIFILM Wako Pure Chemical Corporation, 198-15781), nonessential amino acids (Sigma-Aldrich, M7145) and 20 mM HEPES (Gibco, 15630106). IFN-γ-producing cells were assessed using the BD Mouse IFN-γ ELISPOT Set (BD Biosciences, 551083). Splenocytes (5 × 10□ cells/well) were cocultured for 24 h with γ-irradiated B16F10 cells (10 Gy; 2.5 × 10³ cells/well). Phorbol 12-myristate-13-acetate (PMA, 0.2 ng/mL; Focus Biomolecules, 10-2165) and ionomycin (500 ng/mL; Sigma-Aldrich, I0634) were used as a positive control. Spots were developed with the BD Pharmingen AEC Substrate Kit (BD Biosciences, 551015) and quantified using Fiji.

### Necrosis versus apoptosis assay

Cell-death modality was assessed using the Necrosis vs Apoptosis Assay Kit (ImmunoChemistry Technologies, 9148). B16F10 cells were seeded at 3.0 × 10□ cells/well in 6-well plates 1 day before treatment. Collected cells were stained with FAM-FLICA (1:50) for 60 min at 37 °C and, after washing, with 7-AAD at 5 μg/mL for 20 min at 4 °C in the dark. Fluorescence was measured using an Attune NxT cytometer.

### Western blotting

B16F10 cells were lysed in RIPA buffer containing 1% Triton X-100, complete protease inhibitor cocktail and 10 mM phenylmethylsulfonyl fluoride. To recover proteins from detached/dead cells, culture supernatant was centrifuged and the pellet was added to the lysate. Equal protein amounts were separated by SDS-PAGE and transferred to PVDF membranes. Cleaved caspase-3 (Asp175) (Cell Signaling Technology, 9661, RRID: AB_2341188), total caspase-3 (Cell Signaling Technology, 9662, RRID: AB_331439) and β-actin antibodies (Sigma-Aldrich, A5441, RRID: AB_476744) were used, and signals were detected with Immobilon Western Chemiluminescent HRP Substrate (Merck, WBKLS0100).

### **γ**H2AX detection

B16F10 cells were fixed after treatment with 4% paraformaldehyde, permeabilized with 0.1% Triton X-100, blocked with 5% BSA/PBS and incubated overnight at 4 °C with anti-γH2AX phospho-S139 antibody (Abcam, ab81299, RRID: AB_1640564, clone: EP854(2)Y; 1:500). Alexa Fluor 568 goat anti-rabbit IgG (H+L) (Thermo Fisher Scientific, A11036, RRID: AB_10563566; 1:1000) was used as secondary antibody, followed by Hoechst 33342. Images were acquired with a KEYENCE microscope using a CFI Plan Apo 20× objective (NA 0.75; Nikon).

### Long-term culture

B16F10 cells were seeded at 2.0 × 10□ cells/well in 24-well glass-bottom plates and treated under each condition. For culture longer than 4 days, medium was replaced every 3 or 4 days. Phase-contrast images were acquired with a KEYENCE microscope using a Plan Apo Lambda 20× objective (NA 0.75; Nikon).

### RNA extraction and microarray hybridization

Untreated B16F10 cells and cells exposed to 1 MBq/mL ²¹¹At for 24 h were collected. RNA was extracted using TRIzol Reagent (Invitrogen, 15596026) and 2-propanol precipitation and quantified using a NanoDrop Lite (Thermo Fisher Scientific). RNA (250 ng) was processed with the GeneChip WT PLUS Reagent Kit (Thermo Fisher Scientific, 902280); cDNA/cRNA intermediates were generated, fragmented and labeled, and the hybridization cocktail was loaded onto a Clariom S mouse cartridge (Thermo Fisher Scientific, 902930). Hybridization was performed in a GeneChip Hybridization Oven 645, washing in a GeneChip Fluidics Station 450 (Thermo Fisher Scientific, 00-0079), and scanning in a GeneChip Scanner 3000 7G (Affymetrix, 00-0210).

### DAMP-related measurements

DAMP-related release was assessed in B16F10 and LLC culture supernatants. HMGB1, HSP70/HSPA1A and ATP-related luminescence were measured using the Mouse/Rat HMGB1 ELISA Kit (Arigo Biolaboratories, ARG81310), Human/Mouse/Rat Total HSP70/HSPA1A DuoSet IC ELISA (R&D Systems, DYC1663-2) and CellTiter-Glo 2.0 assay (Promega, G9241), respectively, according to the manufacturers’ protocols. Cell-surface calreticulin was assessed 3 h after treatment by flow cytometry after staining with PE anti-calreticulin antibody (Abcam, ab209577, RRID: AB_2943490, clone: EPR3924; 1:200) and 7-AAD (ImmunoChemistry Technologies, 6163).

### Statistical analysis

Statistical analyses were performed with GraphPad Prism 11. One-way ANOVA followed by Dunnett’s multiple-comparisons test was used when multiple treatment groups were compared with a single control. Two-way ANOVA was used for two-factor experiments; the post hoc procedure is specified in each figure legend. *P* < 0.05 was considered statistically significant. For GSEA, NES and FDR *q*-values are reported.

## Results

### ²¹¹At-damaged B16F10 cells suppress subsequent pulmonary metastasis

To determine whether ²¹¹At-damaged tumor cells influence subsequent tumor progression, we established a cell-vaccination model. B16F10 cells were damaged ex vivo and administered subcutaneously on day 0; viable B16F10 cells were injected intravenously on day 7, and pulmonary metastasis was assessed on day 21 (Fig. 1A). The experiment was conducted under combined immune-checkpoint blockade with anti-PD-1 and anti-CTLA-4 antibodies during priming. The 24-h Trypan Blue signals of cells used for vaccination were broadly comparable across ultraviolet (UV), etoposide (Eto) (21) and ²¹¹At conditions (38.3–43.7% of untreated control; Supplementary Table S1). Residual ²¹¹At activity at administration was approximately 0.92 kBq per mouse. Compared with phosphate-buffered saline (PBS), cells damaged with 1 MBq/mL ²¹¹At markedly reduced pulmonary metastatic nodules, with stronger suppression than the 0.1 MBq/mL ²¹¹At and UV groups (Fig. 1B and C). Eto-damaged cells also suppressed metastasis. These findings indicate that the biological state induced by high-activity ²¹¹At exposure can support antitumor protection in this model.

**Fig. 1.**
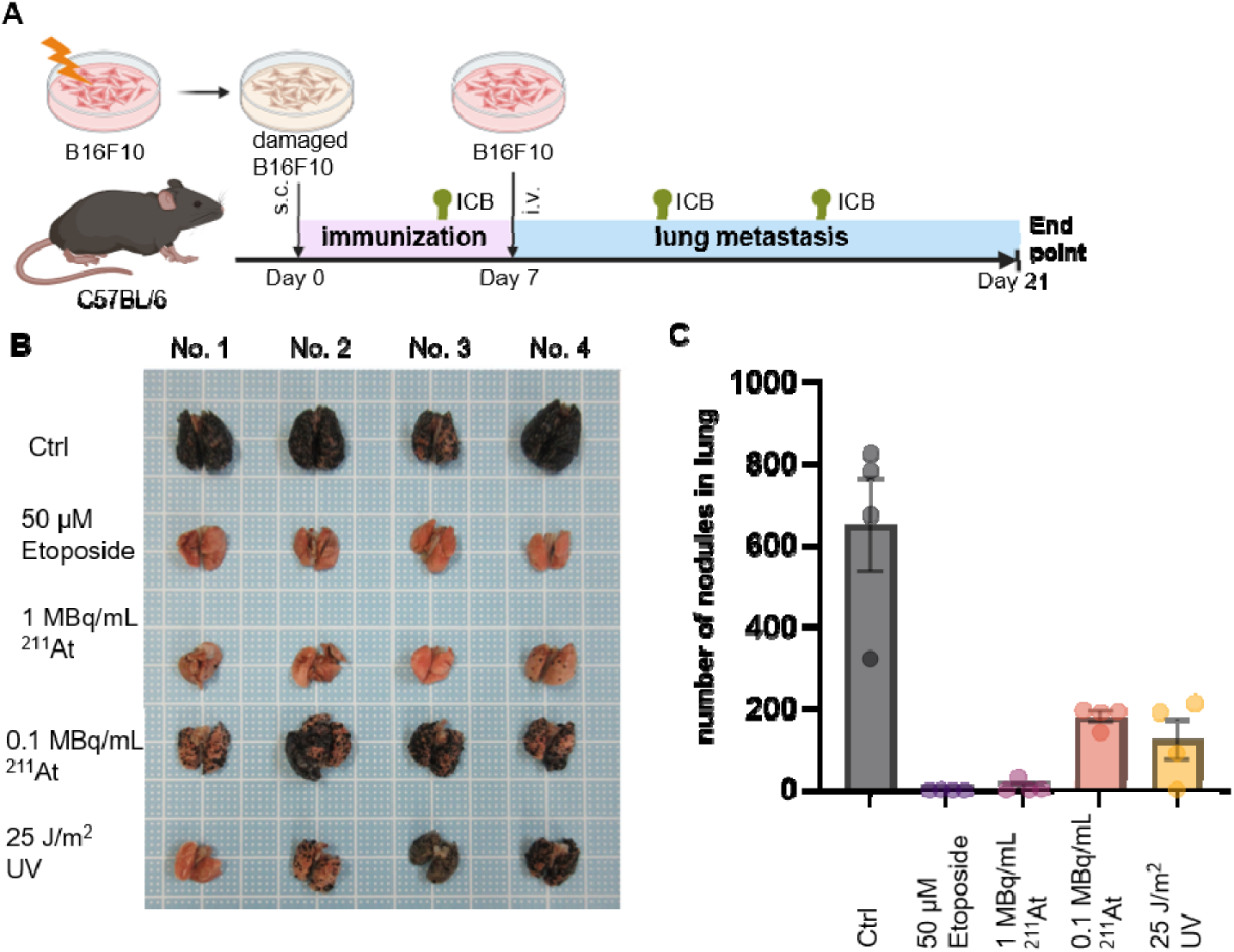
²¹¹At-damaged B16F10 cells suppress subsequent pulmonary metastasis. **A,** Schematic of the lung-metastasis experiment. B16F10 cells were damaged ex vivo with 50 μM etoposide, 25 J/m² UV irradiation, or 0.1 or 1 MBq/mL ²¹¹At and administered subcutaneously on day 0 under immune-checkpoint blockade during priming. Viable B16F10 cells were injected intravenously on day 7, and lungs were analyzed on day 21. **B,** Representative lung images at the endpoint. **C,** Quantification of visible pulmonary nodules. Data are mean ± SEM, n = 4 mice per group. Scheme in (**A**) created in BioRender.

### Damaged tumor cells activate antigen-presenting cells and systemic immune responses

To examine innate immune activation, BMDCs, BMDMs and RAW264.7 cells were cocultured with damaged B16F10 cells or exposed to their culture supernatants (Fig. 2A). Direct coculture with ²¹¹At-damaged B16F10 cells increased CD86 expression, assessed by relative geometric mean fluorescence intensity (gMFI), in an activity-dependent manner (Fig. 2B). Conditioned medium from damaged cells also increased CD86 expression, indicating that soluble factors contribute to antigen-presenting-cell activation in addition to direct cell-cell interactions.

**Fig. 2.**
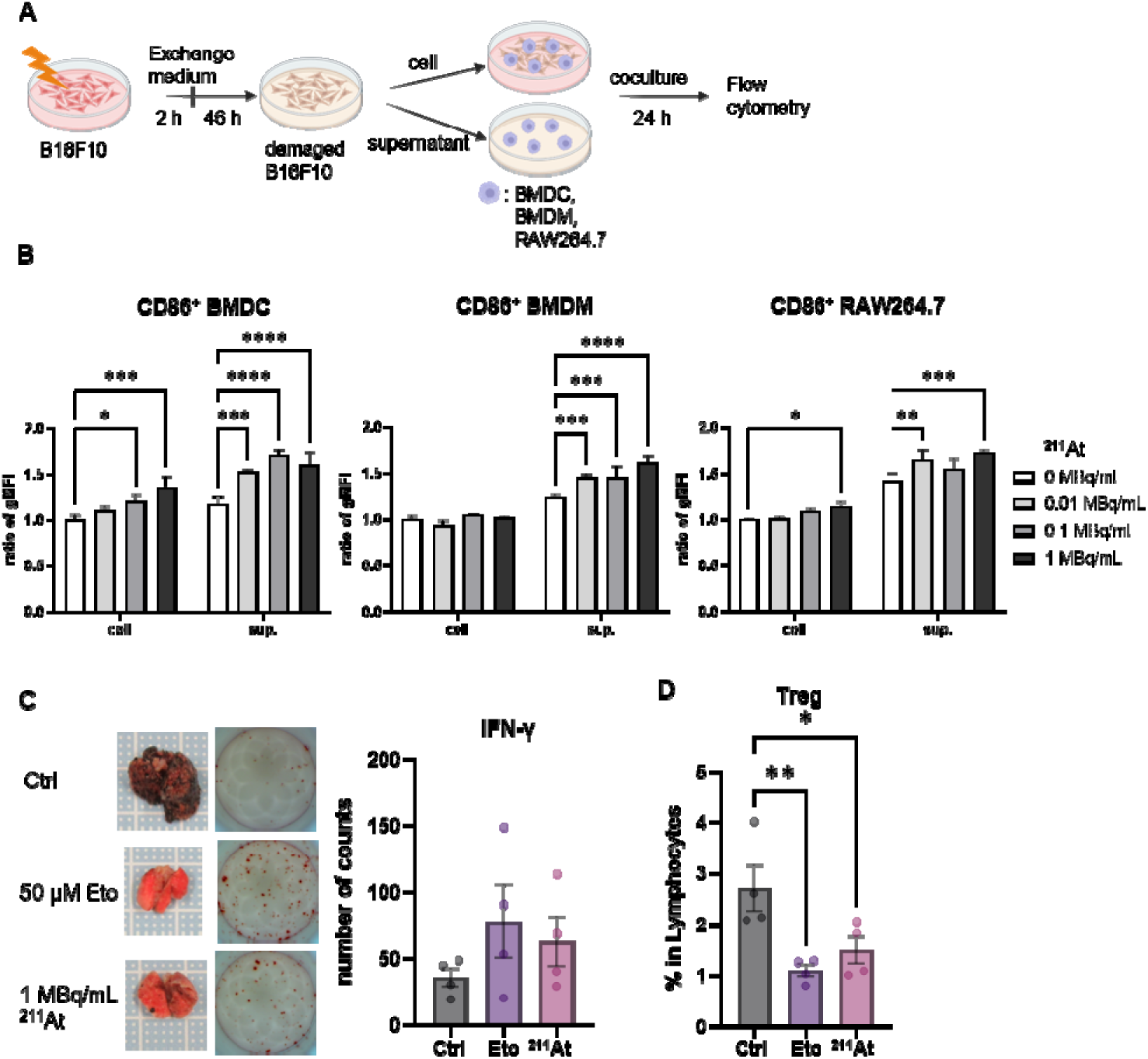
²¹¹At-damaged cells activate BMDCs, BMDMs and RAW264.7 cells and induce systemic immune responses. **A,** Schematic of coculture experiments using damaged B16F10 or LLC cells, their conditioned medium and antigen-presenting cells (APCs). Culture medium was exchanged 2 h after ²¹¹At treatment. **B,** Relative CD86 geometric mean fluorescence intensity (gMFI) in BMDCs, BMDMs and RAW264.7 cells after exposure to damaged tumor cells or conditioned medium. Data are mean ± SD, *n* = 3. \**P* < 0.05, \*\**P* < 0.01, \*\*\**P* < 0.001 and \*\*\*\**P* < 0.0001 by two-way ANOVA followed by Dunnett’s multiple-comparisons test. **C,** Representative lung images and IFN-γ ELISPOT analysis of splenocytes after B16F10 restimulation. **D,** CD4□CD25□FOXP3□ Tregs in splenocytes. Data are mean ± SEM, *n* = 4 mice per group. \**P* < 0.05 and \*\**P* < 0.01 by one-way ANOVA followed by Dunnett’s multiple-comparisons test. Scheme in (**A**) created in BioRender.

At day 21, splenocytes from vaccinated mice produced IFN-γ after B16F10 restimulation in an ELISPOT assay, and the proportion of regulatory T cells (Tregs) was reduced (Fig. 2C and D). Together, these results show that ²¹¹At-damaged B16F10 cells can activate antigen-presenting cells and are associated with a systemic cellular immune response in the pulmonary-metastasis model.

### Vaccination with ²¹¹At-damaged cells induces features of adaptive immunity after a second tumor challenge

We next evaluated immune responses after a second intravenous B16F10 challenge, using Eto-damaged cells as a DNA-damage-associated comparator. Mice received damaged B16F10 cells on day 0, viable B16F10 cells on day 7 and a second intravenous B16F10 injection on day 21, with analysis on day 27 (Fig. 3A). Naïve controls received only the day-21 challenge. Pulmonary metastases were scarcely detectable in mice vaccinated with ²¹¹At-damaged cells and in naïve mice (Fig. 3B). Because naïve mice were observed for only 6 d after their first tumor-cell injection, this comparison primarily informs the immune phenotype after the second challenge rather than providing a stand-alone measure of rechallenge protection.

**Fig. 3.**
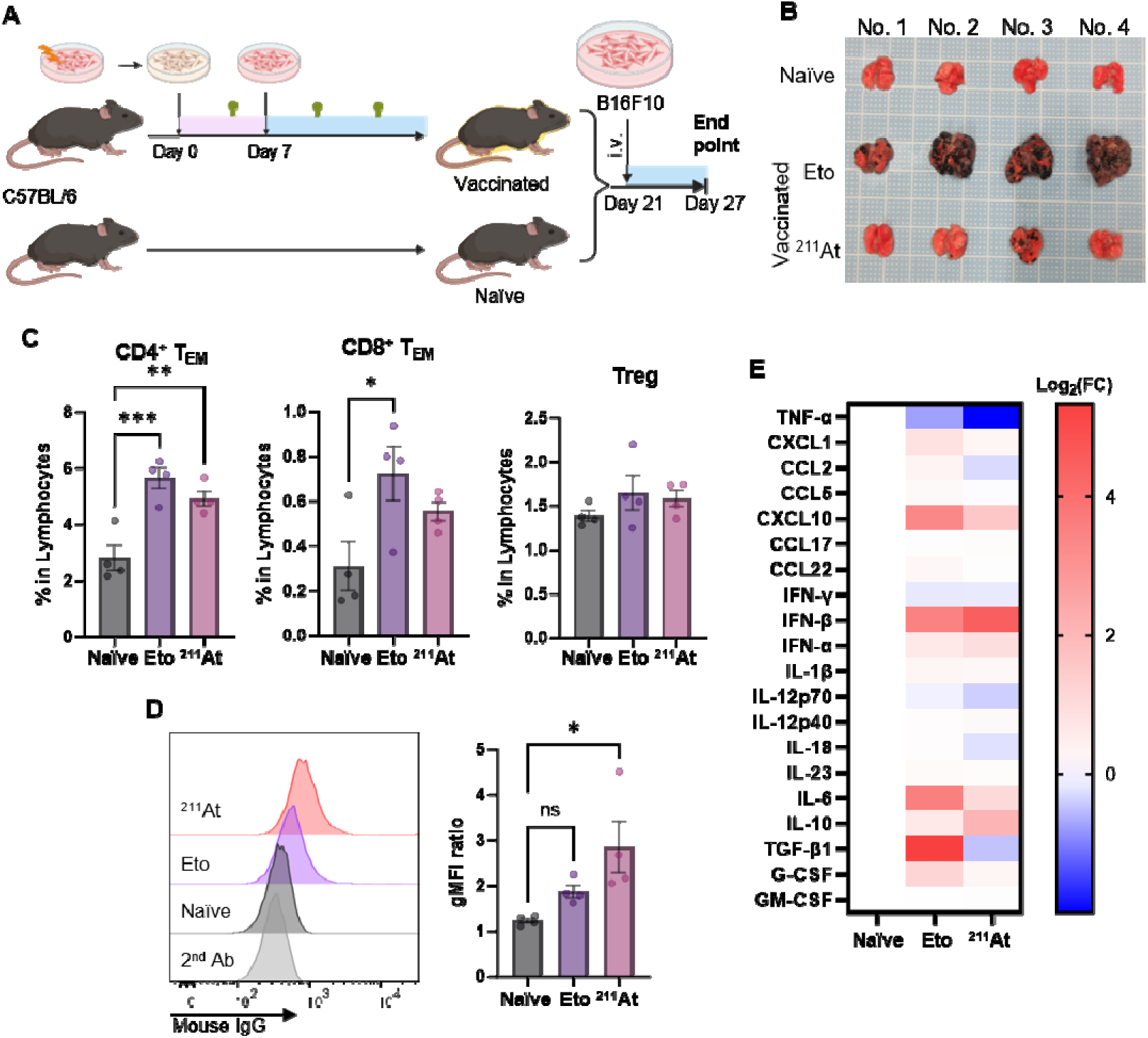
Vaccination with ²¹¹At-damaged cells induces features of systemic adaptive immunity after a second tumor challenge. **A,** Schematic of the second-challenge experiment. ²¹¹At- or Eto-damaged B16F10 cells were used for vaccination; Eto-damaged cells served as a DNA-damage-associated comparator. Vaccinated mice and naïve controls received intravenous B16F10 cells on day 21 and were analyzed on day 27. **B,** Representative lung images. **C,** CD4_J or CD8_J effector-memory T cells (T_EM_) and Tregs. **D,** B16F10-binding mouse IgG histograms and geometric mean fluorescence intensity ratio. **E,** Plasma cytokine/chemokine heatmap shown as log2 fold change. Data are mean ± SEM, *n* = 4 mice per group. \**P* < 0.05 and \*\**P* < 0.01 by one-way ANOVA followed by Dunnett’s multiple-comparisons test. Scheme in (**A**) created in BioRender.

Mice vaccinated with damaged cells showed increased CD4□ and CD8□ effector-memory T cells (T_EM_) without an increase in Treg proportion relative to naïve mice (Fig. 3C). B16F10-binding IgG was detected in plasma and was greater in the ²¹¹At-damaged group than in the Eto-damaged group (Fig. 3D). Plasma profiling showed type I IFN- and CXCL10-related changes after the second challenge, whereas TGF-β was lower in the ²¹¹At-damaged group and higher in the Eto-damaged group (Fig. 3E). These associations do not establish whether TGF-β differences were a cause or consequence of metastatic burden. Depletion of CD4□, CD8□ or CD122□ cells increased pulmonary metastasis to varying degrees (Supplementary Fig. S2), supporting a contribution of lymphocyte-mediated immunity.

### ²¹¹At induces persistent DNA damage, membrane-intact growth arrest and senescence-like changes without prominent SASP secretion at high activity

We next characterized the tumor-cell state associated with immunostimulatory activity. WST-8 assays showed marked reductions in relative metabolic/proliferative signal after ²¹¹At or anticancer-drug treatment, whereas LDH release was substantially lower after ²¹¹At than after Eto or doxorubicin (DOX) (Fig. 4A). Thus, the reduced WST-8 signal after ²¹¹At should not be interpreted simply as the proportion of dead cells; it is consistent with strong suppression of proliferation/metabolic activity in cells with relatively preserved membrane integrity. Flow cytometry showed no prominent acute apoptosis or necrosis under the ²¹¹At conditions (Supplementary Fig. S3), and caspase-3 cleavage was limited compared with Eto (Supplementary Fig. S4).

**Fig. 4.**
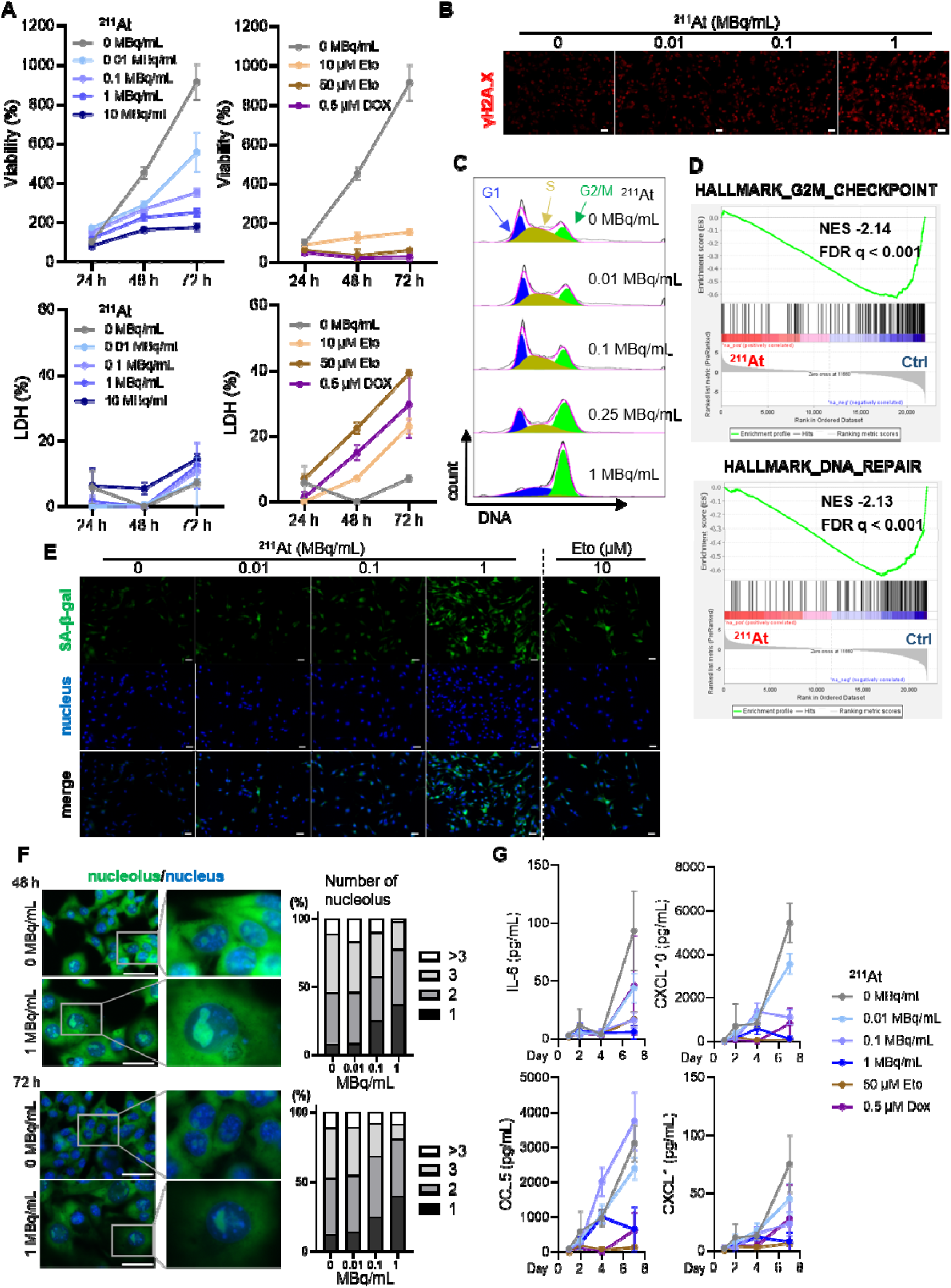
²¹¹At induces persistent DNA damage, membrane-intact growth arrest and senescence-like features without prominent SASP secretion at 1 MBq/mL. **A,** Relative WST-8 signal (viability) and LDH release 24–72 h after treatment. *n* = 3 for biological replicates. **B,** γH2AX immunofluorescence after ²¹¹At treatment. Scale bar, 50 μm. **C,** DNA-content profiles showing G1, S and G2/M distribution. **D,** Selected GSEA results from B16F10 cells exposed to 1 MBq/mL ²¹¹At for 24 h versus untreated cells. Positive NES values indicate enrichment in ²¹¹At-treated cells; negative NES values indicate enrichment in untreated cells. Gene sets with FDR *q*-values reported as 0.000 by GSEA are shown as FDR *q* < 0.001. **E,** SA-β-gal staining after ²¹¹At or Eto treatment. Scale bar, 50 μm. **F,** Nucleus/nucleolus images and distribution of nucleoli per nucleus at 48 and 72 h. (48 h: *n* = 186, 138, 83, 100; 72 h: *n* = 155, 168, 64, 58. n represents the number of nuclei analyzed in a single experiment.) Scale bar, 50 μm. **G,** Time-course analysis of bulk concentrations of SASP-related cytokines/chemokines, including IL-6, CXCL10, CCL5 and CXCL1, in culture supernatants after ²¹¹At or drug treatment. *n* = 3 for biological replicates. NES, normalized enrichment score; FDR, false-discovery rate. Data are mean ± SD. \*\**P* < 0.01 and \*\*\*\**P* < 0.0001 by one-way ANOVA followed by Dunnett’s multiple-comparisons test.

²¹¹At increased γH2AX and induced G2/M accumulation (Fig. 4B and C). GSEA (27, 28) showed strong negative enrichment of proliferation- and cell-cycle-related programs, including the HALLMARK_G2M_CHECKPOINT gene set (NES, −2.14; FDR *q* < 0.001), and suppression of HALLMARK_DNA_REPAIR (NES, −2.13; FDR *q* < 0.001) (Fig. 4D; Supplementary Table S2). Residual B16F10 cells became enlarged and flattened during prolonged culture, with persistent suppression of colony-like growth (Supplementary Fig. S5). Senescence-associated β-galactosidase increased (Fig. 4E; Supplementary Fig. S6), and nucleolar analysis showed enlarged nucleoli and fewer nucleoli per nucleus (Fig. 4F). These findings support a persistent, senescence-like injury state rather than simple acute apoptosis or necrosis.

GSEA also showed enrichment of inflammatory and chemokine-related programs, including HALLMARK_INFLAMMATORY_RESPONSE (NES, 1.52; FDR *q* = 0.032), whereas translation, mRNA splicing, glycolysis and mTORC1-related programs were suppressed (Supplementary Table S2). Mature SASP-associated gene sets were not positively enriched at 24 h. Consistently, bulk supernatant concentrations of IL-6, CXCL10, CCL5 and CXCL1 did not show a marked increase in the 1 MBq/mL ²¹¹At group (Fig. 4G). Because secretion was not normalized to viable cell number or biomass, these data are interpreted as absence of a prominent increase in bulk SASP-factor concentrations rather than proof of unchanged per-cell secretion.

### ²¹¹At induces cytosolic dsDNA without uniformly increasing classical DAMP release

ROS increased 30 min after ²¹¹At treatment (Fig. 5A). By contrast, HMGB1, HSP70, ATP and cell-surface calreticulin did not show consistent activity-dependent increases in B16F10 or LLC cells (Supplementary Fig. S7). LLC cells also showed a different pattern of APC activation (Supplementary Fig. S8), supporting tumor-cell-context dependence. Activation by conditioned medium is therefore unlikely to be explained by a single classical DAMP and may involve multiple soluble factors, extracellular vesicles or nucleic-acid-related signals.

**Fig. 5.**
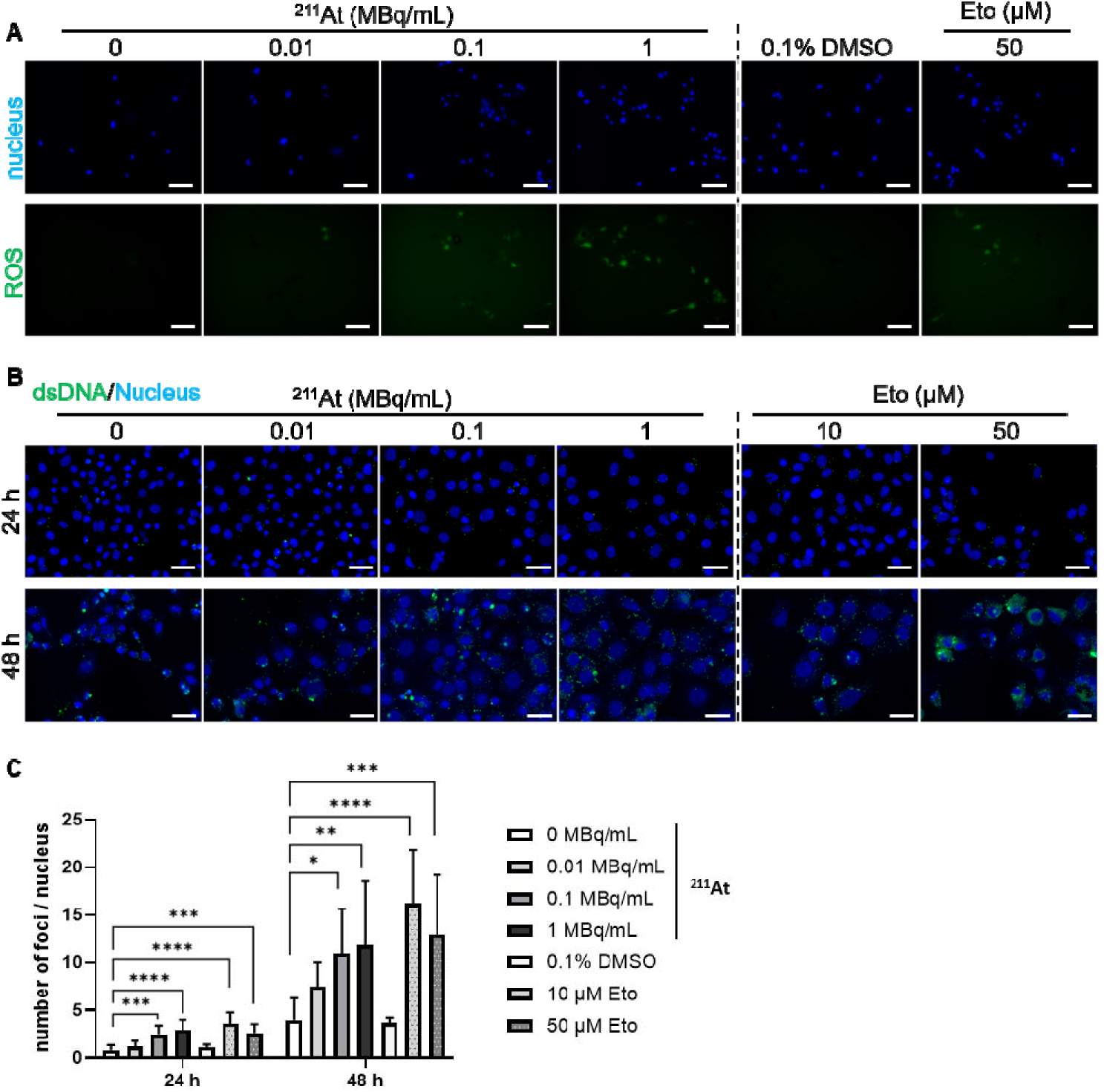
²¹¹At increases early ROS and cytosolic dsDNA. **A,** ROS imaging 30 min after treatment; nuclei are shown in blue and ROS in green. Scale bar, 100 μm. **B,** Cytosolic dsDNA at 24 and 48 h after treatment; nuclei are shown in blue and dsDNA in green. Scale bar, 50 μm. **C,** Quantification of cytosolic dsDNA foci per nucleus. The numbers of microscopic fields of view analyzed were *n* = 10, 6, 8, 11, 8, 10, and 10 at 24 h and *n* = 9, 12, 11, 9, 8, 9 and 8 at 48 h, respectively. Data are mean ± SD. \**P* < 0.05, \*\**P* < 0.01, \*\*\**P* < 0.001 and \*\*\*\**P* < 0.0001 by one-way ANOVA followed by Dunnett’s multiple-comparisons test.

Cytosolic dsDNA foci increased in an activity-dependent manner after ²¹¹At injury, were detectable at 24 h and were more prominent at 48 h (Fig. 5B and C). Together with γH2AX, G2/M accumulation and suppression of DNA-repair and replication programs, these findings identify persistent DNA damage with cytosolic dsDNA accumulation as a prominent feature of the ²¹¹At-damaged-cell state.

### STING inhibition attenuates the immunostimulatory effect of ²¹¹At-damaged cells

Because cytosolic dsDNA lies upstream of cGAS–STING signaling, we examined the effect of the murine STING inhibitor C-176 (Fig. 6A). Vaccination with B16F10 cells damaged by 1 MBq/mL ²¹¹At suppressed pulmonary metastasis, whereas C-176 increased pulmonary nodule counts in these mice (Fig. 6B). The CD4□ central-memory T-cell (T_CM_) analysis in Fig. 6C was performed after pooling blood by treatment group and is therefore interpreted descriptively rather than as an individual-mouse inferential analysis. C-176 also reduced the B16F10-binding IgG response (Fig. 6D). Overall, the data support a contribution of STING signaling to immune activation induced by ²¹¹At-damaged cells.

**Fig. 6.**
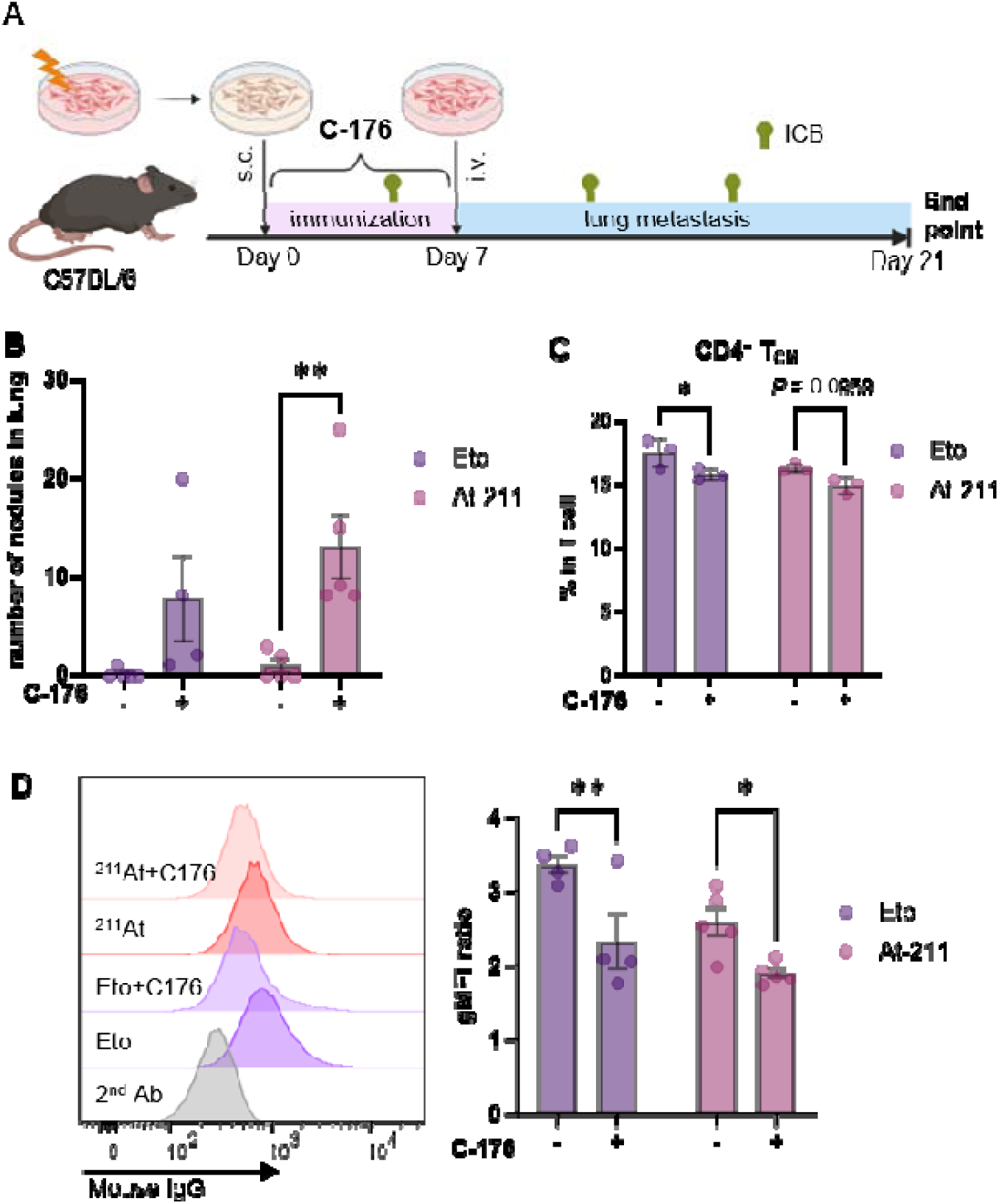
STING inhibition attenuates immunity induced by ²¹¹At-damaged cells. **A,** Schematic of the STING-inhibition experiment with C-176 administered intraperitoneally at 750 nmol in 200 μL corn oil daily from day 0 to day 7. Group sizes: Eto, *n* = 5; Eto + C-176, *n* = 4; ²¹¹At, *n* = 5; ²¹¹At + C-176, *n* = 5. **B,** Pulmonary nodule counts. **C,** Circulating CD4_J central-memory T-cell (T_CM_) values. Blood from individual mice was pooled by treatment group before analysis; the three plotted measurements therefore do not represent independent mice and this panel is interpreted descriptively. Data are mean ± SD. **D,** B16F10-binding mouse IgG histograms and geometric mean fluorescence intensity. Because the blood of one mouse in the Eto-treated group underwent hemolysis and plasma could not be collected, *n* = 4. Data in panels (**B** and **D**) are mean ± SEM. \**P* < 0.05 and \*\**P* < 0.01 by two-way ANOVA followed by Šídák multiple-comparisons test. Scheme in (**A**) created in BioRender.

## Discussion

This study shows that ²¹¹At-damaged B16F10 melanoma cells can function as an immunostimulatory substrate in immunocompetent mice. The findings extend the radiobiology of targeted alpha therapy beyond direct cytotoxicity by linking persistent DNA damage and cytosolic dsDNA with IFN- and chemokine-related transcriptional remodeling, antigen-presenting-cell activation and STING-associated antitumor immunity.

The immune effect was not explained simply by the amount of acute cell death. Cells exposed to ²¹¹At and UV had similar 24-h Trypan Blue signals, yet the 1 MBq/mL ²¹¹At group produced stronger metastasis suppression. For subsequent, more detailed assessment of cellular viability, we used the WST-8 assay; importantly, however, WST-8 reflects metabolic and proliferative activity rather than directly measuring the fraction of live cells. ²¹¹At caused a strong loss of WST-8 signal with comparatively limited LDH release and caspase-3 cleavage, together with persistent γH2AX, G2/M accumulation, suppression of DNA-replication and DNA-repair programs, enlarged and flattened morphology, SA-β-gal positivity, and nucleolar alterations. Because mature SASP programs and bulk SASP-factor concentrations were not prominently increased at 1 MBq/mL, we interpret this as a DNA-damage-associated, senescence-like and biosynthetically suppressed injury state rather than classical senescence with a mature SASP.

The GSEA results strengthen this interpretation. Negative enrichment of DNA-repair, replication, translation and metabolic programs is consistent with a persistent injury state rather than a transient repair response, whereas enrichment of type I IFN, inflammatory and chemokine-related signatures provides a plausible molecular bridge to innate and adaptive immune activation. GSEA alone, however, does not demonstrate STING activation within B16F10 cells.

Comparisons with UV- or Eto-induced states should remain cautious because we did not perform matched transcriptomic profiling of these comparators. Published studies show that UV injury and chemotherapy-induced senescence can each involve DNA-damage, inflammatory and cell-cycle programs, with substantial cell-type- and time-dependent heterogeneity (31–33). The 24-h ²¹¹At signature should therefore be viewed as an early high-LET injury state observed under the present conditions rather than a uniquely ²¹¹At-specific transcriptional program.

The DAMP results also argue against explaining the phenotype solely as classical immunogenic cell death. Calreticulin exposure, ATP, HMGB1 and HSP release are established hallmarks (17–20), yet these readouts did not increase uniformly in B16F10 and LLC cells. Instead, ROS, persistent DNA damage, cytosolic dsDNA and IFN/chemokine-related transcriptional changes were more consistent features. Irradiated cells can transfer immunostimulatory DNA through extracellular vesicles (34), providing one potential mechanism for the conditioned-medium activity observed here.

This interpretation does not imply that ²¹¹At or radiation generally fails to induce DAMPs. ²¹¹At-based treatment has produced apoptosis together with DNA damage in other tumor models (35), and external-beam irradiation can induce calreticulin exposure, HMGB1 or ATP release in a dose-, time- and tumor-cell-dependent manner (36–38). Our data instead indicate that, in B16F10 and LLC cells under the present conditions, persistent DNA damage and cytosolic dsDNA accumulation were more prominent than uniform release of classical DAMPs.

Pharmacological STING inhibition provides functional support for the nucleic-acid-sensing hypothesis (13, 14, 22, 23, 34, 39). C-176 attenuated metastasis suppression and the B16F10-binding IgG response. Systemic C-176 cannot distinguish tumor-cell-intrinsic STING from STING in host immune cells, and recent studies support potential cooperation between these compartments: DNA-damaged cells can engage intrinsic and extrinsic STING signaling after phagocytosis, and cDC1 STING–IRF3 signaling can license antitumor immunity (40, 41). The present data therefore support STING involvement without assigning the effect to a specific cellular compartment.

Adaptive immune findings were also multifaceted. After the second tumor challenge, mice vaccinated with ²¹¹At-damaged cells showed increased effector-memory T cells and B16F10-binding IgG, and lymphocyte depletion reduced protection. The higher TGF-β concentration in the Eto-damaged group was associated with greater metastatic burden, but the experiment cannot determine direction of causality. These results are consistent with a model in which damaged tumor cells provide both tumor antigens and DNA-damage-associated inflammatory signals.

For targeted alpha therapy, an important implication is that the immunologically favorable exposure may not necessarily coincide with the exposure that maximizes immediate cell death. A window may exist in which tumor cells undergo durable proliferative arrest and cytosolic-DNA accumulation while retaining sufficient structural integrity to participate in immune activation. Radionuclide, target antigen, internalization, proximity to the nucleus, intratumoral distribution, activity and dose rate are all likely to influence this balance.

This study has several limitations. The ex vivo vaccination model does not reproduce the spatial and temporal heterogeneity of absorbed dose during in vivo targeted radiopharmaceutical therapy, and ICIs were used during priming. We did not calculate cellular or nuclear absorbed dose, intracellular localization, decays per cell or dose rate, and the contribution of the approximately 0.92 kBq residual activity administered with damaged cells cannot be quantitatively excluded. The STING experiment used systemic pharmacological inhibition rather than genetic models. In addition, blood was pooled by treatment group for the CD4□ central-memory analysis in Fig. 6C; this panel therefore does not provide individual-mouse biological replication.

We also did not perform matched GSEA for UV- or Eto-damaged cells, and SASP-factor concentrations were not normalized to viable cell number or biomass. Future studies should combine absorbed-dose assessment with time-resolved transcriptomics, genetic dissection of tumor- and host-cell STING, individual-animal immune phenotyping and fractionation of soluble, extracellular-vesicle and nucleic-acid components. Such studies will be required to establish how broadly this injury state applies across radiopharmaceuticals and tumor types.

In conclusion, ²¹¹At-damaged B16F10 cells acquired a proliferation-incompetent, DNA-damage-associated state characterized by cytosolic dsDNA accumulation, IFN- and chemokine-related transcriptional remodeling and STING-associated immune activation. The response was not dominated by uniform release of classical DAMPs or a mature SASP. These findings support treatment designs that consider immunogenic conversion of tumor cells alongside direct alpha-particle cytotoxicity.

## Data Availability

The data generated in this study are available within the article and its Supplementary Data files. Additional data are available from the corresponding author upon reasonable request. The microarray data generated during this study are available in the Gene Expression Omnibus database under accession number GSE341281.

## Acknowledgments

This work was supported by the Japan Society for the Promotion of Science (K. Kabayama: JP21H02080, JP25K02604, A. Shimoyama: JP25K01914, K. Fukase: JP25H00006, JP20H05675); the Japan Agency for Medical Research and Development (AMED; A. Shimoyama: JP223fa727001s0301); the Fukushima Institute for Research, Education and Innovation (K. Kabayama: JPFR24040302, JPFR25040302, JPFR26040302); Fusion Oriented Research for disruptive Science and Technology from the Japan Science and Technology Agency (JST; A. Shimoyama: JPMJFR230C); the Japan Initiative for World-leading Vaccine Research and Development Centers; and the Center for Advanced Modalities and DDS (CAMaD; K. Fukase). We would like to thank Prof. F. Yu (Tongji University, School of Medicine, China) for providing the LLC cell line; and Dr. H. Kanoh (Keio University, Institute for Advanced Biosciences, Japan) for providing the RAW264.7 cell line. The authors used Claude (Anthropic) for English language editing. The authors reviewed all edits and take full responsibility for the content.

## Authors’ Contributions

**K. Mayusumi:** Conceptualization, investigation, methodology, writing–original draft, writing–review and editing. **S. Magara:** Investigation, methodology, writing–review and editing. **X. Huang:** Writing–review and editing. **K. Kaneda-Nakashima:** Methodology. **K. Ooe:** Resources. **M. Murakami:** Resources. **A. Toyoshima:** Resources. **H. Haba:** Resources. **X. Yin:** Resources. **K. Fukase:** Conceptualization, investigation, supervision, funding acquisition, writing–review and editing. **A. Shimoyama:** Supervision, funding acquisition. **K. Kabayama:** Conceptualization, investigation, methodology, supervision, funding acquisition, writing–review and editing.

## References

1. Sgouros G, Roeske JC, McDevitt MR, Palm S, Allen BJ, Fisher DR, et al. MIRD Pamphlet No. 22 (Abridged): Radiobiology and Dosimetry of α-Particle Emitters for Targeted Radionuclide Therapy. J Nucl Med 2010;51:311–28.

2. Targeted Alpha Therapy Working Group; Parker C, Lewington V, Shore N, Kratochwil C, Levy M, Lindén O, et al. Targeted Alpha Therapy, an Emerging Class of Cancer Agents: A Review. JAMA Oncol 2018;4:1765–72.

3. Pouget JP, Constanzo J. Revisiting the Radiobiology of Targeted Alpha Therapy. Front Med (Lausanne) 2021;8:692436.

4. Parker C, Nilsson S, Heinrich D, Helle SI, O’Sullivan JM, Fosså SD, et al. Alpha Emitter Radium-223 and Survival in Metastatic Prostate Cancer. N Engl J Med 2013;369:213–23.

5. Kratochwil C, Bruchertseifer F, Giesel FL, Weis M, Verburg FA, Mottaghy F, et al. 225Ac-PSMA-617 for PSMA-Targeted α-Radiation Therapy of Metastatic Castration-Resistant Prostate Cancer. J Nucl Med 2016;57:1941–44.

6. Zalutsky MR, Vaidyanathan G. Astatine-211-labeled radiotherapeutics: an emerging approach to targeted alpha-particle radiotherapy. Curr Pharm Des 2000;6:1433–55.

7. Guérard F, Gestin JF, Brechbiel MW. Production of [211At]-astatinated radiopharmaceuticals and applications in targeted alpha-particle therapy. Cancer Biother Radiopharm 2013;28:1–20.

8. Kaneda-Nakashima K, Zhang Z, Manabe Y, Shimoyama A, Kabayama K, Watabe T, et al. α-Emitting cancer therapy using ²¹¹At-AAMT targeting LAT1. Cancer Sci 2021;112:1132–40.

9. Zalutsky MR, Reardon DA, Akabani G, Coleman RE, Friedman AH, Friedman HS, et al. Clinical Experience with α-Particle–Emitting ²¹¹At: Treatment of Recurrent Brain Tumor Patients with ²¹¹At-Labeled Chimeric Antitenascin Monoclonal Antibody 81C6. J Nucl Med 2008;49:30–38.

10. Andersson H, Cederkrantz E, Bäck T, Divgi C, Elgqvist J, Himmelman J, et al. Intraperitoneal α-Particle Radioimmunotherapy of Ovarian Cancer Patients: Pharmacokinetics and Dosimetry of ²¹¹At-MX35 F(ab′)_2_—A Phase I Study. J Nucl Med 2009;50:1153–60.

11. Watabe T, Hatano K, Naka S, Sasaki H, Kamiya T, Shirakami Y, et al. First-in-human SPECT/CT imaging of [211At]PSMA-5: targeted alpha therapy in a patient with refractory prostate cancer. Eur J Nucl Med Mol Imaging 2025;52:2253–55.

12. Watabe T, Mukai K, Naka S, Sasaki H, Kamiya T, Hayakawa T, et al. First-in-Human Study of [²¹¹At]NaAt as Targeted α-Therapy in Patients with Radioiodine-Refractory Thyroid Cancer (Alpha-T1 Trial). J Nucl Med 2025;66:1941–47.

13. Deng L, Liang H, Xu M, Yang X, Burnette B, Arina A, et al. STING-Dependent Cytosolic DNA Sensing Promotes Radiation-Induced Type I Interferon-Dependent Antitumor Immunity in Immunogenic Tumors. Immunity 2014;41:843–52.

14. Vanpouille-Box C, Alard A, Aryankalayil MJ, Sarfraz Y, Diamond JM, Schneider RJ, et al. DNA exonuclease TREX1 regulates radiotherapy-induced tumour immunogenicity. Nat Commun 2017;8:15618.

15. Gorin JB, Ménager J, Gouard S, Maurel C, Guilloux Y, Faivre-Chauvet A, et al. Antitumor Immunity Induced after α Irradiation. Neoplasia 2014;16:319–28.

16. Li M, Liu D, Lee D, Cheng Y, Baumhover NJ, Marks BM, et al. Targeted Alpha-Particle Radiotherapy and Immune Checkpoint Inhibitors Induces Cooperative Inhibition on Tumor Growth of Malignant Melanoma. Cancers (Basel) 2021;13:3676.

17. Galluzzi L, Vitale I, Warren S, Adjemian S, Agostinis P, Buqué Martinez A, et al. Consensus guidelines for the definition, detection and interpretation of immunogenic cell death. J Immunother Cancer 2020;8:e000337.

18. Obeid M, Tesniere A, Ghiringhelli F, Fimia GM, Apetoh L, Perfettini JL, et al. Calreticulin exposure dictates the immunogenicity of cancer cell death. Nat Med 2007;13:54–61.

19. Apetoh L, Ghiringhelli F, Tesniere A, Obeid M, Ortiz C, Criollo A, et al. Toll-like receptor 4-dependent contribution of the immune system to anticancer chemotherapy and radiotherapy. Nat Med 2007;13:1050–59.

20. Michaud M, Martins I, Sukkurwala AQ, Adjemian S, Ma Y, Pellegatti P, et al. Autophagy-dependent anticancer immune responses induced by chemotherapeutic agents in mice. Science 2011;334:1573–77.

21. Sriram G, Milling LE, Chen JK, Kong YW, Joughin BA, Abraham W, et al. The injury response to DNA damage in live tumor cells promotes antitumor immunity. Sci Signal 2021;14:eabc4764.

22. Glück S, Guey B, Gulen MF, Wolter K, Kang TW, Schmacke NA, et al. Innate immune sensing of cytosolic chromatin fragments through cGAS promotes senescence. Nat Cell Biol 2017;19:1061–70.

23. Dou Z, Ghosh K, Vizioli MG, Zhu J, Sen P, Wangensteen KJ, et al. Cytoplasmic chromatin triggers inflammation in senescence and cancer. Nature 2017;550:402–6.

24. Sato N, Yano S, Toyoshima A, Haba H, Komori Y, Shibata S, et al. Development of a production technology of ²¹¹At at the RIKEN AVF cyclotron: (i) production of ²¹¹At in the ²□□Bi(α,2n)²¹¹At reaction. RIKEN Accel Prog Rep 2017;50:262.

25. Shirakami Y, Watabe T, Obata H, Kaneda K, Ooe K, Teramoto T, et al. Synthesis of [²¹¹At]4-astato-L-phenylalanine by dihydroxyboryl-astatine substitution reaction in aqueous solution. Sci Rep 2021;11:12982.

26. Ito K, Manabe Y, Ohshima S, Maeki M, Tokeshi M, Inaba H, et al. Programmable Antigen-Specific Immunity via Self-Adjuvanting Nanovaccines Co-Delivering Immune Modulators. Angew Chem Int Ed 2026;65:e202520474.

27. Subramanian A, Tamayo P, Mootha VK, Mukherjee S, Ebert BL, Gillette MA, et al. Gene set enrichment analysis: a knowledge-based approach for interpreting genome-wide expression profiles. Proc Natl Acad Sci U S A 2005;102:15545–50.

28. Liberzon A, Birger C, Thorvaldsdóttir H, Ghandi M, Mesirov JP, Tamayo P. The Molecular Signatures Database (MSigDB) hallmark gene set collection. Cell Syst 2015;1:417–25.

29. Spada S, Yamazaki T, Vanpouille-Box C. Detection and quantification of cytosolic DNA. Methods Enzymol 2019;629:17–33.

30. Schindelin J, Arganda-Carreras I, Frise E, Kaynig V, Longair M, Pietzsch T, et al. Fiji: an open-source platform for biological-image analysis. Nat Methods 2012;9:676–82.

31. Shen Y, Kim AL, Du R, Liu L. Transcriptome Analysis Identifies the Dysregulation of Ultraviolet Target Genes in Human Skin Cancers. PLoS One 2016;11:e0163054.

32. Jochems F, Thijssen B, De Conti G, Jansen R, Pogacar Z, Groot K, et al. The Cancer SENESCopedia: a delineation of cancer cell senescence. Cell Rep 2021;36:109441.

33. Fernandez B, Passanisi VJ, Ashraf HM, Spencer SL. Single-cell RNA sequencing reveals a quiescence-senescence continuum and distinct senotypes following chemotherapy. Nat Commun 2026;17:169.

34. Diamond JM, Vanpouille-Box C, Spada S, Rudqvist NP, Chapman JR, Ueberheide BM, et al. Exosomes Shuttle TREX1-Sensitive IFN-Stimulatory dsDNA from Irradiated Cancer Cells to DCs. Cancer Immunol Res 2018;6:910–20.

35. Ma H, Ye T, Qu G, Qin Y, Liao J, Yang Y, et al. Locoregional radionuclide therapy of glioblastoma with [²¹¹At]At-PDA-FAPI. Sci Rep 2025;15:18248.

36. Kesar U, Markelc B, Cemazar M, Sersa G, Jesenko T, Strojan P. Early effects of ionizing radiation on molecular modifications and hallmarks of immunogenic cell death in mouse tumor cells. BMC Cancer 2026;26:401.

37. Gameiro SR, Jammeh ML, Wattenberg MM, Tsang KY, Ferrone S, Hodge JW. Radiation-induced immunogenic modulation of tumor enhances antigen processing and calreticulin exposure, resulting in enhanced T-cell killing. Oncotarget 2014;5:403–16.

38. Golden EB, Frances D, Pellicciotta I, Demaria S, Barcellos-Hoff MH, Formenti SC. Radiation fosters dose-dependent and chemotherapy-induced immunogenic cell death. Oncoimmunology 2014;3:e28518.

39. Haag SM, Gulen MF, Reymond L, Gibelin A, Abrami L, Decout A, et al. Targeting STING with covalent small-molecule inhibitors. Nature 2018;559:269–73.

40. Park S, Ahn J, Barber GN. Antigen-presenting cell activation requires intrinsic and extrinsic STING signalling after the phagocytosis of DNA-damaged cells. Sci Immunol 2024;9:eadk7812.

41. Wang J, Li S, Wang M, Wang X, Chen S, Sun Z, et al. STING licensing of type I dendritic cells potentiates antitumor immunity. Sci Immunol 2024;9:eadj3945.

