## Supplemental Fig S1 for "Astatine-211 injury induces an immunostimulatory melanoma-cell state and promotes systemic antitumor immunity"

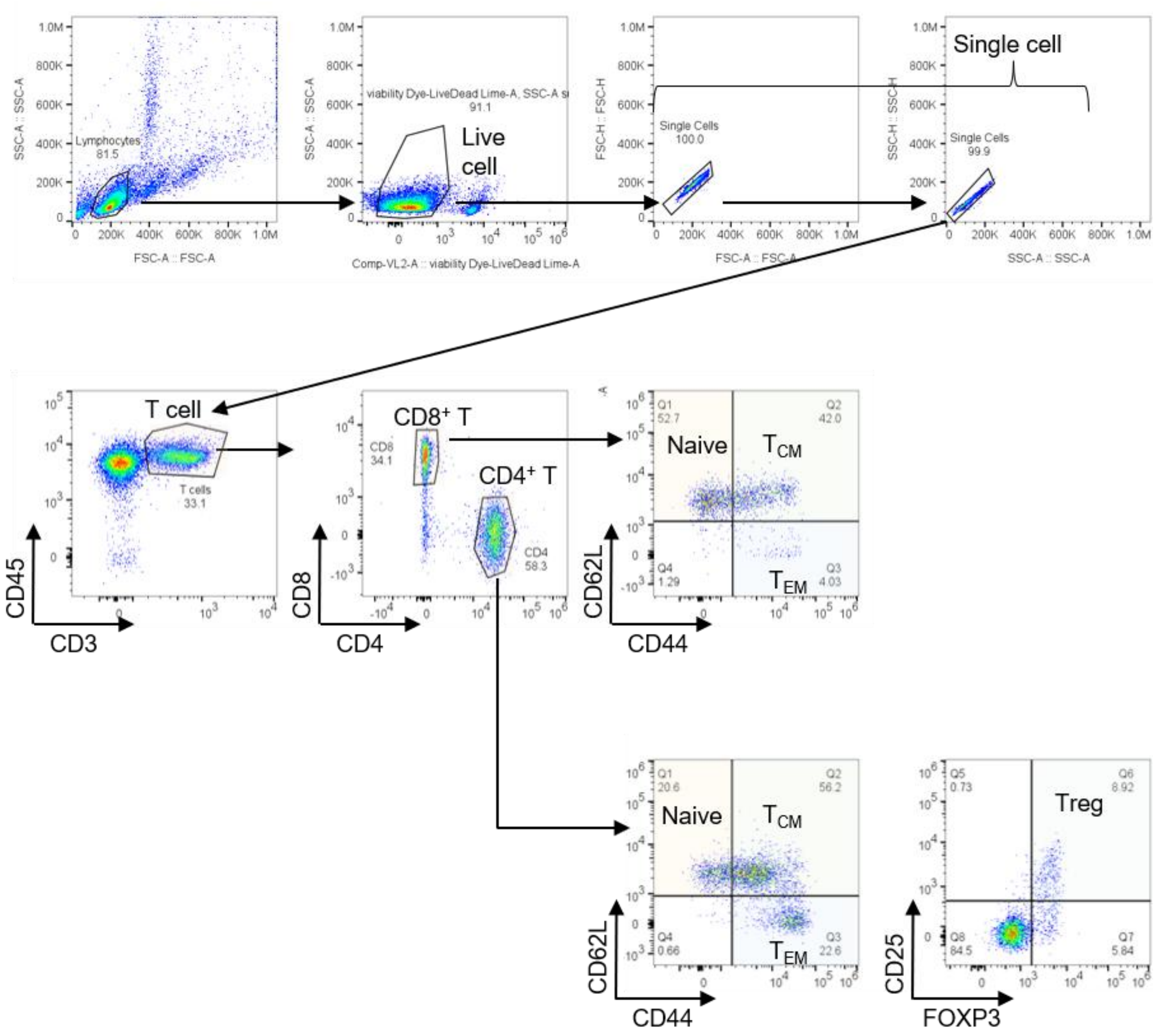


**Supplementary Fig. S1** Flow-cytometric gating strategy for T-cell-subset analysis. Representative plots show sequential gating of lymphocytes, live cells and singlets, followed by identification of $CD45^{+}CD3^{+}$ T cells. $CD4^{+}$ and $CD8^{+}$ T cells were classified as naïve ($CD44^{-}CD62L^{+}$), central-memory (TCM; $CD44^{+}CD62L^{+}$) or effector-memory (TEM; $CD44^{+}CD62L^{-}$) cells. Regulatory T cells (Tregs) were identified within the $CD4^{+}$ T-cell population as $CD25^{+}FOXP3^{+}$ cells.
