## Supplemental Fig S2 for "Astatine-211 injury induces an immunostimulatory melanoma-cell state and promotes systemic antitumor immunity"

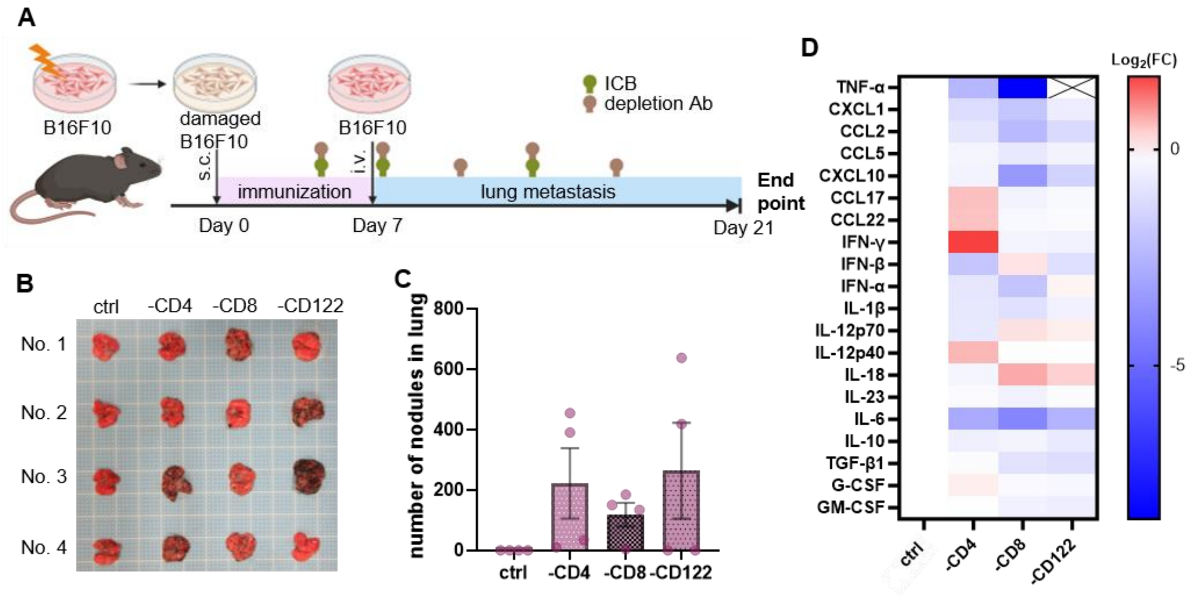


**Supplementary Fig. S2** Lymphocyte depletion attenuates the antimetastatic effect induced by $^{211}$At-damaged cells. **A,** Schematic of the depletion experiment. Mice were vaccinated subcutaneously with $^{211}$At-damaged B16F10 cells on day 0 under immune-checkpoint blockade, challenged intravenously with viable B16F10 cells on day 7, and treated with control or depleting antibodies against CD4, CD8 or CD122; lungs and plasma were analyzed on day 21. **B,** Representative lung images. **C,** Visible pulmonary nodules. Data are mean ± SEM, n = 4 mice per group. **D,** Plasma cytokine/chemokine profiles shown as log2-transformed fold changes. ICB, immune-checkpoint blockade. Scheme in (**A**) created in BioRender.
