## Supplemental Fig S3 for "Astatine-211 injury induces an immunostimulatory melanoma-cell state and promotes systemic antitumor immunity"

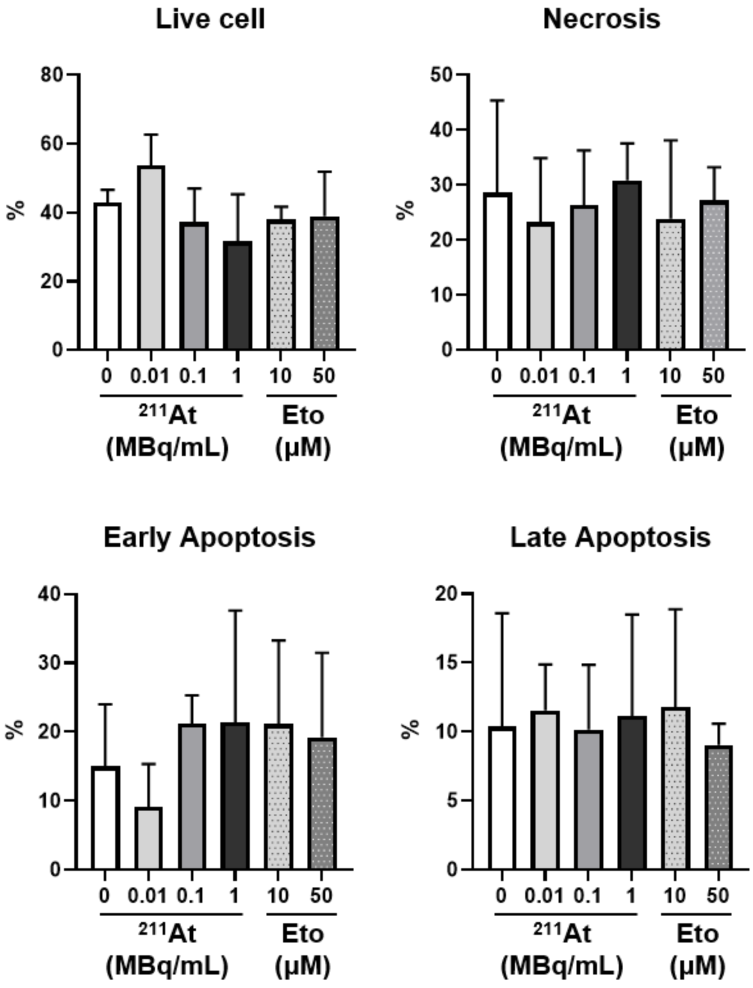


**Supplementary Fig. S3** Apoptosis and necrosis after $^{211}$At or etoposide treatment. Flow-cytometric quantification of live, necrotic, early-apoptotic and late-apoptotic B16F10 cells 24 h after treatment with the indicated $^{211}$At activities or etoposide concentrations. Cell-death states were determined using FAM-FLICA and 7-AAD staining. Data are mean ± SD, *n* = 3.
