## Supplemental Fig S4 for "Astatine-211 injury induces an immunostimulatory melanoma-cell state and promotes systemic antitumor immunity"

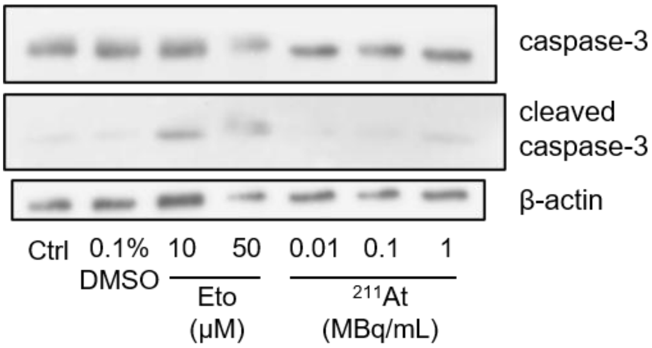


**Supplementary Fig. S4** Caspase-3 processing after $^{211}$At or etoposide treatment. Representative western blots of caspase-3 and cleaved caspase-3 in B16F10 cells 24 h after treatment with the indicated etoposide concentrations or $^{211}$At activities. β-actin was used as the loading control.
