## Supplemental Fig S5 for "Astatine-211 injury induces an immunostimulatory melanoma-cell state and promotes systemic antitumor immunity"

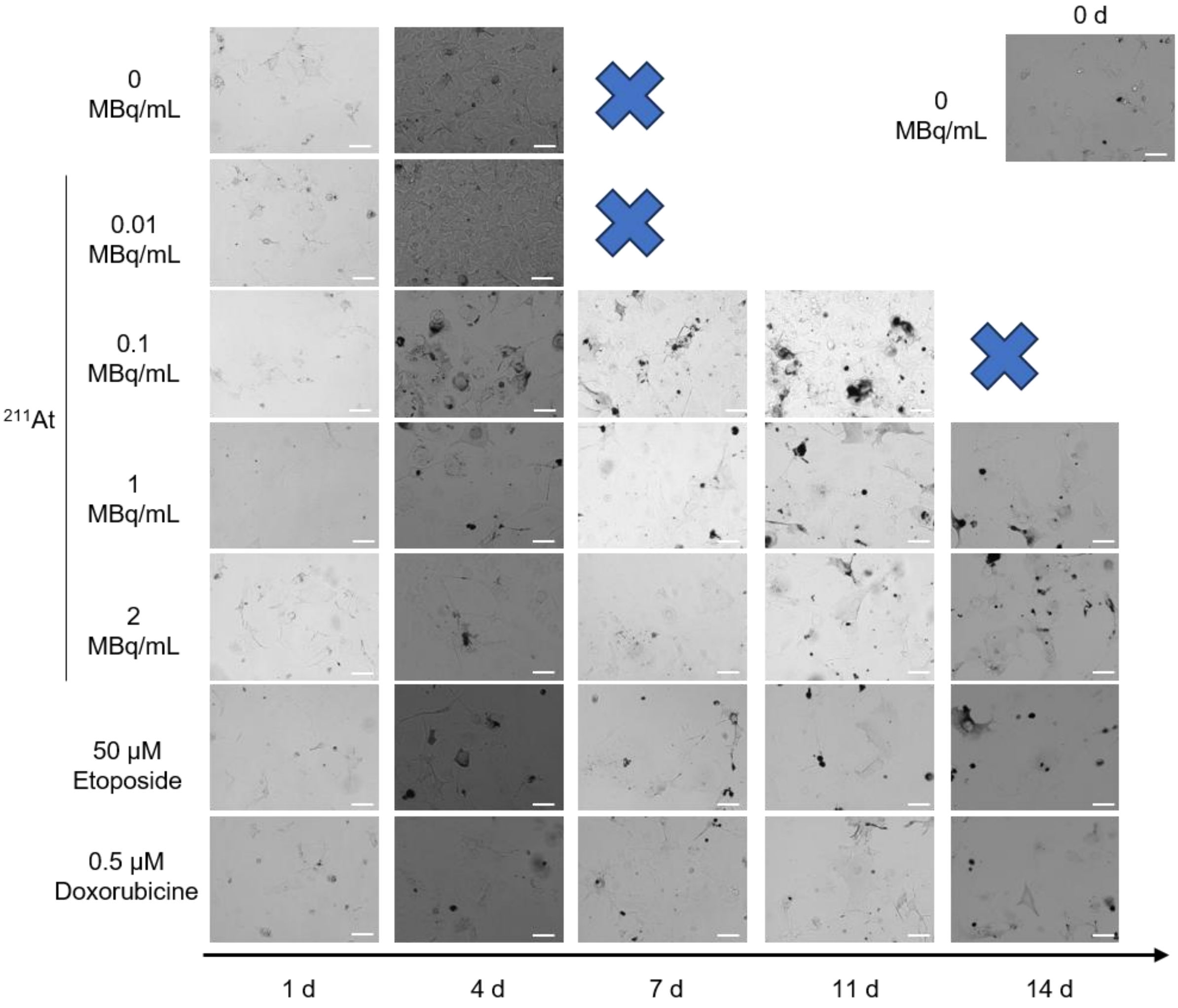


**Supplementary Fig. S5** Long-term morphological changes in B16F10 cells after $^{211}$At or drug treatment. Representative phase-contrast images were acquired 1, 4, 7, 11 and 14 d after treatment with the indicated $^{211}$At activities, 50 µM etoposide or 0.5 µM doxorubicin. Untreated cells at day 0 are shown for reference. Blue crosses indicate time points at which cultures were overgrown. Scale bars, 100 µm.
