## Supplemental Fig S6 for "Astatine-211 injury induces an immunostimulatory melanoma-cell state and promotes systemic antitumor immunity"

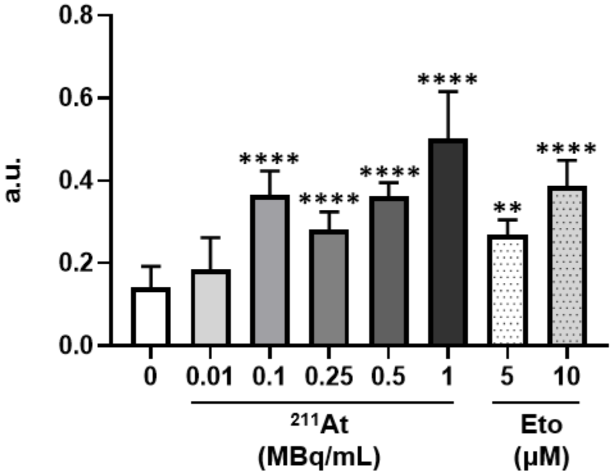


**Supplementary Fig. S6** Increased senescence-associated β-galactosidase activity after $^{211}$At treatment. Senescence-associated β-galactosidase activity in B16F10 cells was quantified 48 h after treatment with the indicated $^{211}$At activities or etoposide concentrations. Data are mean ± SD, $n = 3$. $^{**}P < 0.01$ and $^{****}P < 0.0001$ by one-way ANOVA followed by Dunnett's multiple-comparisons test. a.u., arbitrary units.
