## Supplemental Fig S7 for "Astatine-211 injury induces an immunostimulatory melanoma-cell state and promotes systemic antitumor immunity"

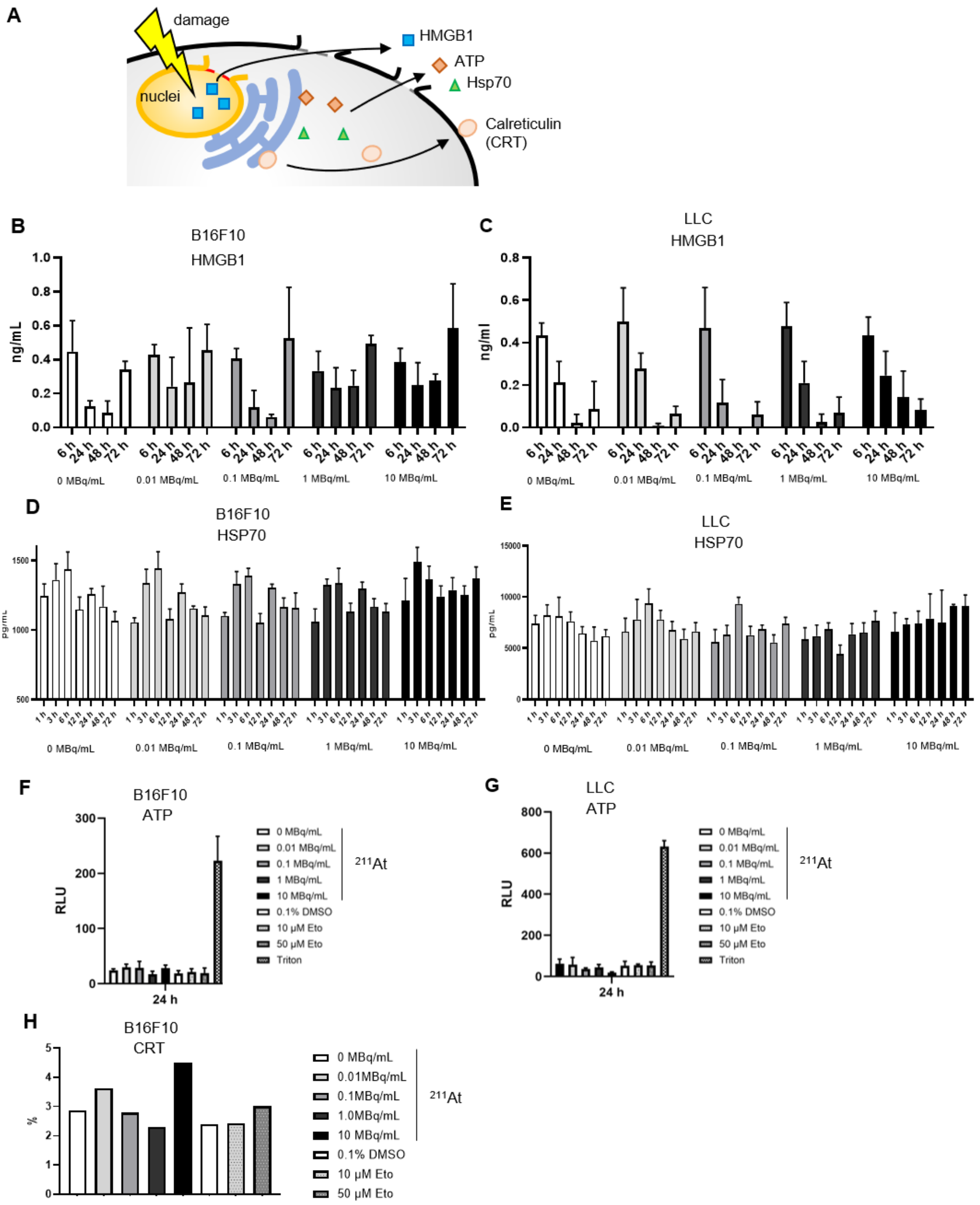


**Supplementary Fig. S7** $^{211}$At does not uniformly increase classical DAMP-related responses. **A,** Schematic of extracellular HMGB1, ATP and HSP70 release and cell-surface calreticulin (CRT) exposure after tumor-cell damage. **B** and **C,** HMGB1 concentrations in culture supernatants from B16F10 (**B**) and LLC (**C**) cells collected 6–72 h after treatment with the indicated $^{211}$At activities. **D** and **E,** HSP70 concentrations in culture supernatants from B16F10 (**D**) and LLC (**E**) cells collected 1–72 h after treatment. **F** and **G**, ATP-related luminescence from B16F10 (**F**) and LLC (**G**) cells 24 h after $^{211}$At or etoposide treatment; Triton X-100 was used as a positive control. **H,** Percentage of cell-surface CRT-positive B16F10 cells 3 h after $^{211}$At or etoposide treatment, measured by flow cytometry. Data in (**B–G**) are mean ± SD, $n = 3$. RLU, relative luminescence units.
