## Supplemental Fig S8 for "Astatine-211 injury induces an immunostimulatory melanoma-cell state and promotes systemic antitumor immunity"

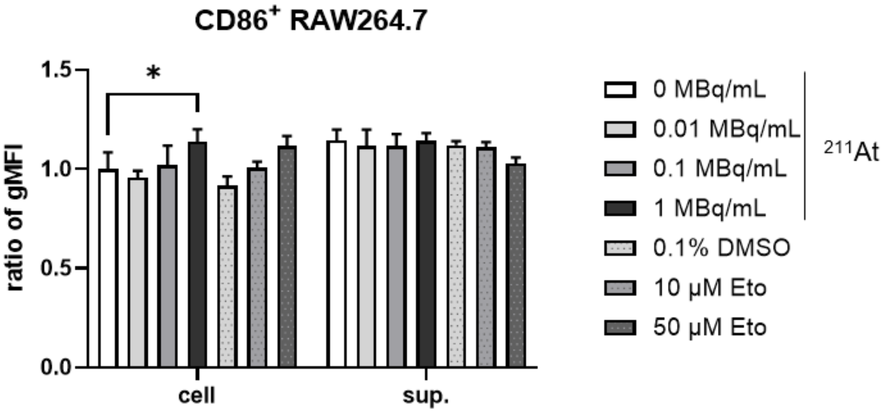


**Supplementary Fig. S8** Relative CD86 geometric mean fluorescence intensity in RAW264.7 cells after exposure to damaged LLC cells or LLC-cell conditioned medium. Data are mean ± SD, $n = 3$. *$P < 0.05$ by two-way ANOVA followed by Dunnett's multiple-comparisons test.
