## Supplemental Table S1 for "Astatine-211 injury induces an immunostimulatory melanoma-cell state and promotes systemic antitumor immunity"

**Supplementary Table S1** Relative Trypan Blue signal of B16F10 cells used for vaccination 24 h after treatment. Cells were treated with 50 µM etoposide, 25 J/m² ultraviolet (UV) irradiation, or 0.1 or 1 MBq/mL $^{211}At$. Values are expressed relative to the untreated control.

| **Treatment** | **Relative Trypan Blue signal at 24 h (% of untreated control)** |
|---|---|
| 50 µM Etoposide | 43.7 |
| 1 MBq/mL $^{211}At$ | 43.6 |
| 0.1 MBq/mL $^{211}At$ | 42.4 |
| 25 J/m² UV | 38.3 |
